# Integrase-RNA interactions underscore the critical role of integrase in HIV-1 virion morphogenesis

**DOI:** 10.1101/2019.12.18.881649

**Authors:** Jennifer Elliott, Jenna E. Eschbach, Pratibha C. Koneru, Wen Li, Maritza Puray Chavez, Dana Townsend, Dana Lawson, Alan N. Engelman, Mamuka Kvaratskhelia, Sebla B. Kutluay

## Abstract

A large number of HIV-1 integrase (IN) alterations, referred to as class II substitutions, exhibit pleotropic effects during virus replication. However, the underlying mechanism for the class II phenotype is not known. Here we demonstrate that all tested class II IN substitutions compromised IN-RNA binding in virions by one of three distinct mechanisms: i) markedly reducing IN levels thus precluding formation of IN complexes with viral RNA; ii) adversely affecting functional IN multimerization and consequently impairing IN binding to viral RNA; iii) directly compromising IN-RNA interactions without substantially affecting IN levels or functional IN multimerization. Inhibition of IN-RNA interactions resulted in mislocalization of the viral ribonucleoprotein complexes outside the capsid lattice, which led to premature degradation of the viral genome and IN in target cells. Collectively, our studies uncover causal mechanisms for the class II phenotype and highlight an essential role of IN-RNA interactions for accurate virion maturation.

## INTRODUCTION

Infectious HIV-1 virions are formed in a multistep process coordinated by interactions between the HIV-1 Gag and Gag-Pol polyproteins, and the viral RNA (vRNA) genome. At the plasma membrane of an infected cell, Gag and Gag-Pol molecules assemble around a vRNA dimer and bud from the cell as a spherical immature virion, in which the Gag proteins are radially arranged [1–3]. As the immature virion buds, the viral protease enzyme is activated and cleaves Gag and Gag-Pol into their constituent domains triggering virion maturation [1, 2]. During maturation the cleaved nucleocapsid (NC) domain of Gag condenses with the RNA genome and *pol*-encoded viral enzymes [reverse transcriptase (RT) and integrase (IN)] inside the conical capsid lattice, composed of the cleaved capsid (CA) protein, which together form the core [1–3].

After infection of a target cell, RT in the confines of the reverse transcription complex (RTC) synthesizes linear double stranded DNA from vRNA [4]. The vDNA is subsequently imported into the nucleus, where the IN enzyme catalyzes its insertion into the host cell chromosome [5, 6]. Integration is mediated by the intasome nucleoprotein complex that consists of a multimer of IN engaging both ends of linear vDNA [7]. While the number of IN protomers required for intasome function varies across Retroviridae, single particle cryogenic electron microscopy (cryo-EM) structures of HIV-1 and Maedi-visna virus indicate that lentivirus integration proceeds via respective higher-order dodecamer and hexadecamer IN arrangements [8, 9], though a lower-order intasome comprised of an HIV-1 IN tetramer was also resolvable by cryo-EM [9].

A number of IN substitutions which specifically arrest HIV-1 replication at the integration step have been described [10]. These substitutions are grouped into class I to delineate them from a variety of other IN substitutions, which exhibit pleiotropic effects and are collectively referred to as class II substitutions [10–12]. Class II IN substitutions or deletion of entire IN impair proper particle assembly [11, 13–25], morphogenesis [11, 15, 21–23, 26–28] and reverse transcription in target cells [10, 11, 17, 19–21, 23, 25–44], in some cases without impacting IN catalytic function [15, 16, 19, 20, 30, 31, 34, 36, 45–47]. A hallmark morphological defect of these viruses is the formation of aberrant viral particles with viral ribonucleoprotein (vRNP) complexes mislocalized outside of the conical CA lattice [11, 15, 21–23, 26–28]. Strikingly similar morphological defects are observed in virions produced from cells treated with allosteric integrase inhibitors (ALLINIs, also known as LEDGINs, NCINIs, INLAIs or MINIs) [26, 27, 48-55]. ALLINIs induce aberrant IN multimerization in virions by engaging the V-shaped pocket at the IN dimer interface, which also provides a principal binding site for the host integration targeting cofactor lens epithelium-derived growth factor (LEDGF)/p75 [50, 54, 56–60]. The recent discovery that HIV-1 IN binds to the vRNA genome in virions and that inhibiting IN-RNA interactions leads to the formation of eccentric particles provided initial clues about the role of IN during virion morphogenesis [28].

HIV-1 IN consists of three independently folded protein domains: the N-terminal domain (NTD), catalytic core domain (CCD), and C-terminal domain (CTD) [7, 61], and vRNA binding is mediated by a constellation of basic residues within the CTD [28]. However, class II IN substitutions are located throughout the entire length of the IN protein [10, 12], which raises the question as to how these substitutions impair virus maturation. The structural basis for IN binding to RNA is not yet known; however, in vitro evidence indicates that IN binds RNA as lower-order multimers, and conversely RNA binding may prevent the formation of higher order IN multimers [28]. Notably, aberrant IN multimerization underlies the inhibition of IN-RNA interactions by ALLINIs [28] and subsequent defects in virion maturation [26-28, 48, 49, 51-55]. Therefore, it seems plausible that class II IN substitutions may exert their effect on virus replication by adversely affecting functional IN multimerization. However, a systematical evaluation of the effects of IN substitutions on IN multimerization, IN-RNA binding, and virion morphology is lacking. As such, it remains an open question how functional IN multimerization and/or IN-RNA interactions influence correct virion morphogenesis.

Eccentric virions generated via class II IN substitutions or ALLINI treatment are defective for reverse transcription in target cells [10, 11, 17, 19–21, 23, 25–44, 48, 49, 51, 54, 58, 62] despite containing equivalent levels of RT and vRNA genome as wild type (WT) particles [26, 63]. In addition, neither the condensation of the viral genome by NC [26, 63] nor its priming [63] appear to be affected. We and others have recently shown that premature loss of the viral genome and IN, as well as spatial separation of RT from vRNPs, may underlie the reverse transcription defect observed in eccentric viruses generated in the presence of ALLINIs or the class II IN R269A/K273A substitutions [59, 64]. These findings support a model in which the capsid lattice or IN binding to vRNA itself is necessary to protect viral components from the host environment upon entering a target cell. Whether the premature loss of the viral genome and IN is a universal outcome of other class II IN substitutions is unknown.

In this work, we aimed to determine the molecular basis of how class II IN substitutions exert their effects on HIV-1 replication. In particular, by detailed characterization of how class II substitutions impact IN multimerization, IN-RNA interactions and virion morphology, we aimed to dissect whether loss of IN binding to vRNA or aberrant IN multimerization underlies the pleiotropic defects observed in viruses bearing class II IN mutations. Remarkably, we found that class II substitutions either prevented IN binding to the vRNA genome or precluded the formation of IN-vRNA complexes through reducing or eliminating IN from virions. We show that IN tetramers have a strikingly higher affinity towards vRNA than IN monomers or dimers, and a large number of class II IN substitutions inhibited IN binding to RNA indirectly through modulating functional IN tetramerization. In contrast, R262A/R263A and R269A/K273A substitutions within the CTD and the K34A change within the NTD did not perturb IN tetramer formation, and thus likely directly interfered with IN binding to RNA. Irrespective of how IN-RNA binding was inhibited, all class II IN mutant viruses formed eccentric particles with vRNPs mislocalized outside of the CA lattice. Subsequently, this led to premature loss of the vRNA genome as well as IN, and spatial separation of RT and CA from the vRNPs in target cells. Taken together, our findings uncover causal mechanisms for the class II phenotype and highlight the essential role of IN-RNA interactions for the formation of correctly matured virions and vRNP stability in HIV-1-infected cells.

## MATERIALS AND METHODS

### Plasmids

The pNLGP plasmid consisting of the HIV-1_NL4-3_-derived Gag-Pol sequence inserted into the pCR/V1 plasmid backbone [65] and the CCGW vector genome plasmid carrying a GFP reporter under the control of the CMV promoter [66, 67] were previously described. The pLR2P-vprIN plasmid expressing a Vpr-IN fusion protein has also been previously described [68]. Mutations in the IN coding sequence were introduced into both the pNLGP plasmid and the HIV-1_NL4-3_ full-length proviral plasmid (pNL4-3) by overlap extension PCR. Briefly, forward and reverse primers containing IN mutations in the *pol* reading frame were used in PCR reactions with antisense and sense outer primers containing unique restriction endonuclease sites (AgeI-sense, NotI-antisense for NLGP and AgeI-sense, EcoRI-antisense for pNL4-3), respectively. The resulting fragments containing the desired mutations were mixed at 1:1 ratio and overlapped subsequently using the sense and antisense primer pairs. The resulting fragments were digested with the corresponding restriction endonucleases and cloned into pNLGP and pNL4-3 plasmids. IN mutations were introduced into the pLR2P-vprIN plasmid using the QuickChange Site-Directed Mutagenesis kit (Agilent Technologies). Presence of the desired mutations and absence of unwanted secondary changes were verified by Sanger sequencing.

### Cells and viruses

HEK293T cells (ATCC CRL-11268) and HeLa-derived TZM-bl cells (NIH AIDS Reagent Program) were maintained in Dulbecco’s modified Eagle’s medium supplemented with 10% fetal bovine serum. MT-4 cells were maintained in RPMI 1640 medium supplemented with 10% fetal bovine serum. CHO K1-derived pgsA-745 cells (CRL-2242, ATCC) that lack a functional xylosyltransferase enzyme and as a result do not produce glycosaminoglycans were maintained in Dulbecco’s modified Eagle’s / F12 (1:1) media supplemented with 10% fetal bovine serum and 1 mM L-glutamine. Single-cycle GFP reporter viruses pseudotyped with vesicular stomatitis virus G protein (VSV-G) were produced by transfection of HEK293T cells with pNLGP-derived plasmids, the CCGW vector genome carrying GFP, and VSV-G expression plasmid at a ratio of 5:5:1, respectively, using polyethyleneimine (PolySciences, Warrington, PA). Full-length viruses pseudotyped with VSV-G were produced by transfecting HEK293T cells with the pNL4-3-derived plasmids and VSV-G plasmid at a ratio of 4:1 (pNL4-3:VSV-G).

### Immunoblotting

Viral and cell lysates were resuspended in sodium dodecyl sulfate (SDS) sample buffer and separated by electrophoresis on Bolt 4-12% Bis-Tris Plus gels (Life Technologies), blotted onto nitrocellulose membranes and probed overnight at 4°C with the following antibodies in Odyssey Blocking Buffer (LI-COR): mouse monoclonal anti-HIV p24 antibody (183-H12-5C, NIH AIDS reagents), mouse monoclonal anti-HIV integrase antibody [69], rabbit polyclonal anti-HIV integrase antibody raised in-house against Q44-LKGEAMHGQVD-C56 peptide and hence unlikely to be affected by the substitutions introduced into IN in this study, rabbit polyclonal anti-HIV-1 reverse transcriptase antibody (6195, NIH AIDS reagents), rabbit polyclonal anti-Vpr antibody (11836, NIH AIDS Reagents), rabbit polyclonal anti-MA antibody (4811, NIH AIDS Reagents). Membranes were probed with fluorophore-conjugated secondary antibodies (LI-COR) and scanned using a LI-COR Odyssey system. IN and CA levels in virions were quantified using Image Studio software (LI-COR). For analysis of the fates of core components in infected cells, antibody incubations were done using 5% non-fat dry milk. Membranes were probed with HRP-conjugated secondary antibodies and developed using SuperSignal^TM^ West Femto reagent (Thermo-Fisher).

### Analysis of reverse transcription products in infected cells

MT-4 cells were grown in 24-well plates and infected with VSV-G pseudotyped pNL4-3 viruses (either WT or class II IN mutant) at a multiplicity of infection (MOI) of 2 in the presence of polybrene. Six h post-infection cells were collected, pelleted by brief centrifugation, and resuspended in PBS. DNA was extracted from cells using the DNeasy Blood and Tissue Kit (Qiagen) as per kit protocol. Quantity of HIV-1 vDNA was measured by Q-PCR using primers specific for early reverse-transcripts.

### Vpr-IN transcomplementation experiments

A class I IN mutant virus (HIV-1_NL4-3_ IN_D116N_) was trans-complemented with class II mutant IN proteins as described previously [68]. Briefly, HEK293T cells grown in 24-well plates were co-transfected with a derivative of the full-length HIV-1_NL4-3_ proviral plasmid bearing a class I IN substitution (pNL4-3_D116N_), VSV-G, and derivatives of the pLR2P-vprIN plasmid bearing class II IN mutations at a ratio of 6:1:3. Two days post-transfection cell-free virions were collected from cell culture supernatants. Integration capability of the trans-complemented class II IN mutants was tested by infecting MT-4 cells and measuring the yield of progeny virions in cell culture supernatants over a 6-day period as described previously [68]. In brief, MT-4 cells were incubated with virus inoculum in 96 V-bottom well plates for 4 h at 37°C after which the virus inoculum was washed away and replaced with fresh media. Immediately following removal of the virus inoculum and during the six subsequent days the quantity of virions present in the culture supernatant was quantified by measuring RT activity using a Q-PCR-based assay [70].

### CLIP experiments

CLIP experiments were conducted as previously described [28, 71, 72]. Cell-free HIV-1 virions were isolated from transfected HEK293T cells. Briefly, cells in 15-cm cell culture plates were transfected with 30 µg full-length proviral plasmid (pNL4-3) DNA containing the WT sequence or indicated *pol* mutations within the IN coding sequence. Cells were grown in the presence of 4-thiouridine for 16 h prior to virus harvest. Two days post transfection cell culture supernatants were collected and filtered through 0.22 μm filters and pelleted by ultracentrifugation through a 20% sucrose cushion using a Beckman SW32-Ti rotor at 28,000 rpm for 1.5 h at 4°C. Virus pellets were resuspended in phosphate-buffered saline (PBS) and UV-crosslinked. Following lysis in RIPA buffer, IN-RNA complexes were immunoprecipitated using a mouse monoclonal anti-IN antibody [69]. Bound RNA was end-labeled with γ-^32^P-ATP and T4 polynucleotide kinase. The isolated protein-RNA complexes were separated by SDS-PAGE, transferred to nitrocellulose membranes and exposed to autoradiography films to visualize RNA. Lysates and immunoprecipitates were also analyzed by immunoblotting using antibodies against IN.

### IN multimerization in virions

HEK293T cells grown on 10-cm dishes were transfected with 10 µg pNL4-3 plasmid DNA containing the WT sequence or indicated *pol* mutations within IN coding sequence. Two days post-transfection cell-free virions collected from cell culture supernatants were pelleted by ultracentrifugation through a 20% sucrose cushion using a Beckman SW41-Ti rotor at 28,000 rpm for 1.5 h at 4°C. Pelleted virions were resuspended in 1X PBS and treated with ethylene glycol bis(succinimidyl succinate) (EGS) (ThermoFisher Scientific), a membrane permeable crosslinker, at a concentration of 1 mM for 30 min at room temperature. Crosslinking was stopped by addition of SDS sample buffer. Samples were subsequently separated on 3-8% Tris-acetate gels and analyzed by immunoblotting using a mouse monoclonal anti-IN antibody [69].

### Size exclusion chromatography (SEC)

All of the mutations were introduced into a plasmid backbone expressing His_6_ tagged pNL4-3-derived IN by QuikChange site directed mutagenesis kit (Agilent) [60]. His_6_ tagged recombinant pNL4-3 WT and mutant INs were expressed in BL21 (DE3) *E. coli* cells followed by nickel and heparin column purification as described previously [60, 73]. Recombinant WT and mutant INs were analyzed on Superdex 200 10/300 GL column (GE Healthcare) with running buffer containing 20 mM HEPES (pH 7.5), 1 M NaCl, 10% glycerol and 5 mM BME at 0.3 mL/min flow rate. The proteins were diluted to 10 µM with the running buffer and incubated for 1 h at 4°C followed by centrifugation at 10,000g for 10 min. Multimeric form determination was based on the standards including bovine thyroglobulin (670,000 Da), bovine gamma-globulin (158,000 Da), chicken ovalbumin (44,000 Da), horse myoglobin (17,000 Da) and vitamin B12 (1,350 Da).

### Analysis of IN-RNA binding in vitro

Following SEC of IN as above, individual fractions of tetramer, dimer and monomer forms were collected and their binding to TAR RNA was analyzed by an Alpha screen assay as described previously [28]. Briefly, 100 nM His_6_ tagged IN fractions (tetramer, dimer and monomer) were incubated with nickel acceptor beads while increasing concentrations of biotinylated-TAR RNA was incubated with streptavidin donor beads in buffer containing 100 mM NaCl, 1 mM MgCl_2_, 1 mM DTT, 1 mg/mL BSA, 25 mM Tris (pH 7.4). Followed by 2-h incubation at 4°C, they were mixed and the reading was taken after 1 h incubation at 4°C by PerkinElmer Life Sciences Enspire multimode plate reader. The Kd values were calculated using OriginLab software.

### Virus production and transmission electron microscopy

Cell-free HIV-1 virions were isolated from transfected HEK293T cells. Briefly, cells grown in two 15-cm cell culture plates (10^7^ cells per dish) were transfected with 30 μg full-length proviral plasmid (pNL4-3) DNA containing the WT sequence or indicated *pol* mutations within IN coding sequence using PolyJet DNA transfection reagent as recommended by the manufacturer (SignaGen Laboratories). Two days after transfection, cell culture supernatants were filtered through 0.22 μm filters and pelleted by ultracentrifugation using a Beckman SW32-Ti rotor at 26,000 rpm for 2 h at 4°C. Fixative (2.5% glutaraldehyde, 1.25% paraformaldehyde, 0.03% picric acid, 0.1 M sodium cacodylate, pH 7.4) was gently added to resulting pellets, and samples were incubated overnight at 4°C. The following steps were conducted at the Harvard Medical School Electron Microscopy core facility. Samples were washed with 0.1 M sodium cacodylate, pH 7.4 and postfixed with 1% osmium tetroxide /1.5% potassium ferrocyanide for 1 h, washed twice with water, once with maleate buffer (MB), and incubated in 1% uranyl acetate in MB for 1 h. Samples washed twice with water were dehydrated in ethanol by subsequent 10 minute incubations with 50%, 70%, 90%, and then twice with 100%. The samples were then placed in propyleneoxide for 1 h and infiltrated overnight in a 1:1 mixture of propyleneoxide and TAAB Epon (Marivac Canada Inc.). The following day the samples were embedded in TAAB Epon and polymerized at 60 °C for 48 h. Ultrathin sections (about 60 nm) were cut on a Reichert Ultracut-S microtome, transferred to copper grids stained with lead citrate, and examined in a JEOL 1200EX transmission electron microscope with images recorded on an AMT 2k CCD camera. Images were captured at 30,000x magnification, and over 100 viral particles per sample were counted by visual inspection.

### Equilibrium density sedimentation of virion core components in vitro

Equilibrium density sedimentation of virion core components was performed as previously described [64]. Briefly, HEK293T cells grown in 10-cm cell culture plates were transfected with 10 µg pNLGP plasmid DNA containing the WT sequence or indicated *pol* mutations within IN coding sequence. Two days post-transfection cell-free virions collected from cell culture supernatants were pelleted by ultracentrifugation through a 20% sucrose cushion using a Beckman SW41-Ti rotor at 28,000 rpm for 1.5 hr at 4°C. Pelleted viral-like particles were resuspended in PBS and treated with 0.5% Triton X-100 for 2 min at room temperature. Immediately after, samples were layered on top of 30-70% linear sucrose gradients prepared in 1X STE buffer (100 mM NaCl, 10 mM Tris-Cl (pH 8.0), 1 mM EDTA) and ultracentrifuged using a Beckman SW55-Ti rotor at 28,500 rpm for 16 h at 4°C. Fractions (500 µL) collected from the top of the gradients were analyzed for IN, CA, and MA by immunoblotting as detailed above.

### Biochemical analysis of virion core components in infected cells

Biochemical analysis of retroviral cores in infected cells was performed as described previously [74]. Briefly, pgsA-745 cells were infected with VSV-G pseudotyped single cycle GFP-reporter viruses or its derivatives synchronously at 4°C. Following the removal of virus inoculum and extensive washes with PBS, cells were incubated at 37°C for 2 h. To prevent loss of vRNA due to reverse-transcription, cells were infected in the presence of 25 µM nevirapine. Post-nuclear supernatants were separated by ultracentrifugation on 10-50% linear sucrose gradients using a Beckman SW55-Ti rotor at 30,000 rpm for 1 h at 4°C. Ten 500 µl fractions from the top of the gradient were collected, and CA, IN, and vRNA in each fraction were analyzed by either immunoblotting or Q-PCR [74]. A SYBR-Green-based Q-PCR assay [70] was used to determine RT activity in the collected sucrose fractions.

### Visualization of vRNA in infected cells

Viral RNA was visualized in infected cells according to the published multiplex immunofluorescent cell-based detection of DNA, RNA and Protein (MICDDRP) protocol [75]. VSV-G pseudotyped HIV-1_NL4-3_ virus stocks were prepared as described above and concentrated 40X using a lentivirus precipitation solution (ALSTEM). PgsA-745 cells were plated on 1.5 mm collagen-treated coverslips (GG-12-1.5-Collagen, Neuvitro) placed in 24-well plates one day prior to infection. Synchronized infections were performed by incubating pre-chilled virus inoculum on the cells for 30 min at 4°C. Cells were infected with WT virus at a MOI of 0.5, or with an equivalent number (normalized by RNA copy number) of IN mutant viral particles. After removal of the virus inoculum cells were washed with PBS and either immediately fixed with 4% paraformaldehyde, or incubated at 37°C for 2 h before fixing. To prevent loss of vRNA due to reverse-transcription, cells were infected and incubated in the presence of 25 µM nevirapine. Following fixation, cells were dehydrated with ethanol and stored at −20°C. Prior to probing for vRNA, cells were rehydrated, incubated in 0.1% Tween in PBS for 10 min, and mounted on slides. Probing was performed using RNAScope probes and reagents (Advanced Cell Diagnostics). Briefly, coverslips were treated with protease solution for 15 min in a humidified HybEZ oven (Advanced Cell Diagnostics) at 40 °C. The coverslips were then washed with PBS and pre-designed anti-sense probes [75] specific for HIV-1 vRNA were applied and allowed to hybridize with the samples in a humidified HybEZ oven at 40 °C for 2 h. The probes were visualized by hybridizing with preamplifiers, amplifiers, and finally, a fluorescent label. First, pre-amplifier 1 (Amp 1-FL) was hybridized to its cognate probe for 30 min in a humidified HybEZ oven at 40 °C. Samples were then subsequently incubated with Amp 2-FL, Amp 3-FL, and Amp 4A-FL for 15 min, 30 min, and 15 min respectively. Between adding amplifiers, the coverslips were washed with a proprietary wash buffer. Nuclei were stained with DAPI diluted in PBS at room temperature for 5 min. Finally, coverslips were washed in PBST followed by PBS and then mounted on slides using Prolong Gold Antifade.

### Microscopy and image quantification

Images were taken using a Zeiss LSM 880 Airyscan confocal microscope equipped with a ×63/1.4 oil-immersion objective using the Airyscan super-resolution mode. 10 images were taken for each sample using the ×63 objective. Numbers of nuclei and vRNA punctae in images were counted using Volocity software (Quorum Technologies). The number of vRNA punctae per 100 nuclei were recorded at 0 h post-infection (hpi) and 2 hpi for each virus, and the number at 2 hpi compared to the number at 0 hpi.

### Analysis of the fate of vRNA genome in MT4 cells

MT-4 cells were infected with VSV-G pseudotyped HIV-1 NL4-3 WT or equivalent number of mutant viruses (normalized by RT activity) synchronously at 4°C. After removal of virus inoculum and extensive washes with PBS, cells were incubated at 37°C for 6 h in the presence of 25 µM nevirapine. Immediately after synchronization (0 h) and at 2 and 6 h post-infection samples were taken from the infected cultures and RNA was isolated using TRIzol Reagent. The amount of viral genomic RNA was measured by Q-RT-PCR.

## RESULTS

### Characterization of the replication defects of class II IN mutant viruses

Substitutions in IN that exhibited a class II phenotype (i.e. assembly, maturation or reverse transcription defects [10-44, 76, 77] or affected IN multimerization [46, 78–81] were selected from past literature. The location of these substitutions depicted on the cryo-EM structure of the tetrameric HIV-1 intasome complex [9] indicate that many are positioned at or near monomer-monomer or dimer-dimer interfaces (Fig. 1A-B). While not apparent in the tetrameric intasome complex, the CTD mediates IN tetramer-tetramer interactions in the higher-order dodecamer IN structure [9] and has also been shown to mediate IN multimerization in vitro [15].

**Figure 1.**
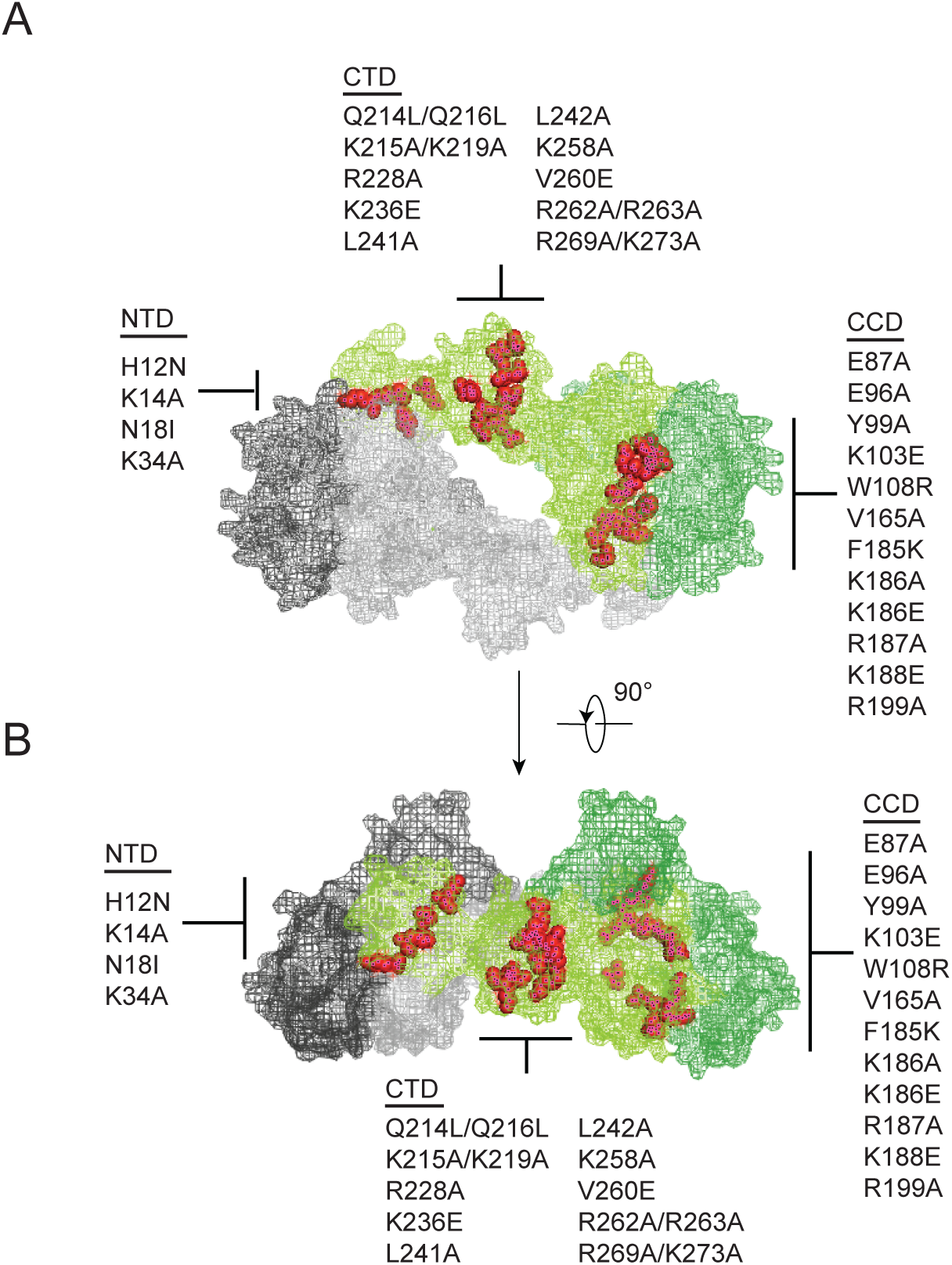
Class II IN substitutions locate throughout IN and cluster at interfaces that mediate IN multimerization. (A) Location of class II IN substitutions used in this study displayed in red on a single IN monomer within the context of the HIV-1 IN tetramer intasome structure consisting of a dimer of dimers (PDB 5U1C). The two dimers are displayed in either gray or green, with individual monomers within each displayed in different shades. The DNA is omitted for clarity. (B) View of the structure displayed in A rotated 90°.

IN mutations were introduced into the replication competent pNL4-3 molecular clone and HEK293T cells were transfected with the resulting plasmids. Cell lysates and cell-free virions were subsequently analyzed for Gag/Gag-Pol expression, processing, particle release and infectivity. While substitutions in IN had no measurable effect on Gag (Pr55) expression, modest effects on Gag processing in cells was visible for several missense mutant viruses including H12N, N18I, K34A, Y99A, K103E, W108R, F185K, Q214L/Q216L, L242A, V260E, as well as the ΔIN mutant (Fig. 2A). Nevertheless, particle release was largely similar between WT and IN mutant viruses, as evident by the similar levels of CA protein present in cell culture supernatants (Fig. 2A, lower panels).

**Figure 2.**
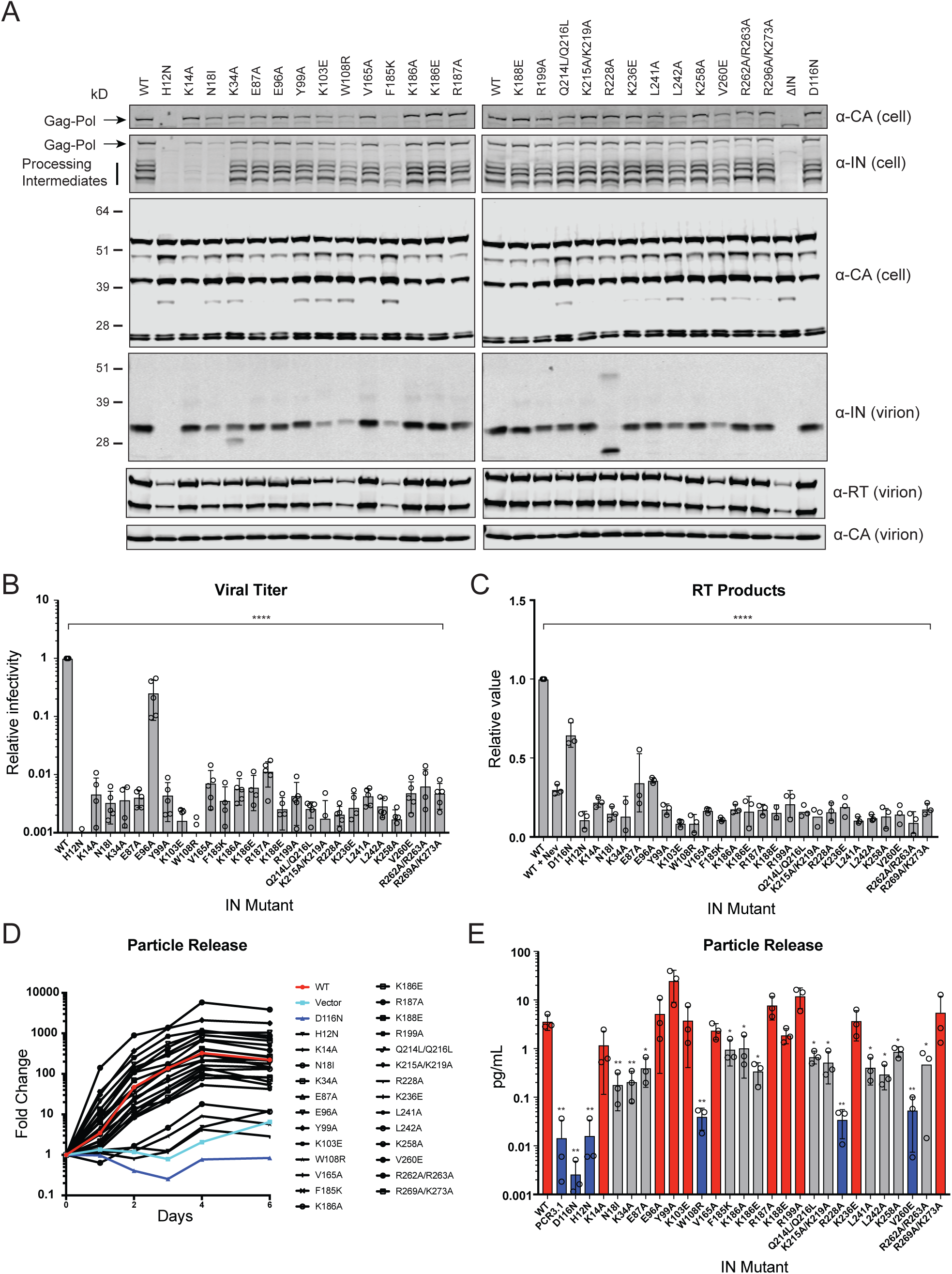
Characterization of the replication defects of class II IN mutant viruses. (A) Immunoblot analysis of Gag and Gag-Pol products in cell lysates and virions. HEK293T cells were transfected with proviral HIV-1_NL4-3_ expression plasmids carrying *pol* mutations encoding for the indicated IN substitutions. Cell lysates and purified virions were harvested two days post transfection and analyzed by immunoblotting for CA, IN and, in the case of virions, RT. Representative image of one of four independent experiments is shown. (B) Infectious titers of WT or IN mutant HIV-1_NL4-3_ viruses in cell culture supernatants were determined on TZM-bl indicator cells. Titer values are expressed relative to WT (set to 1). Columns show average of five independent experiments (open circles) and error bars represent standard deviation (****P < 0.0001, by one-way ANOVA with Dunnett’s multiple comparison test). (C) Relative quantity of reverse-transcribed HIV-1 DNA in MT-4 target cells infected with HIV-1_NL4-3_ at 6 hpi. Quantities of vDNA are expressed relative to WT (set to 1). Columns show average of three independent experiments (open circles) and error bars represent standard deviation (****P < 0.0001, by one-way ANOVA with Dunnett’s multiple comparison test). (D) Representative growth curve of HIV-1_NL4-3_ IN_D116N_ viruses trans-complemented with class II mutant IN proteins in cell culture. Y-axis indicates fold increase in virion yield over day 0 as measured by RT activity in culture supernatants. HIV-1_NL4-3_ IN_D116N_ viruses that were trans-complemented with WT IN, class II mutant INs, IN_D116N_, or an empty vector are denoted as red, black, dark blue and light blue lines respectively. Representative plot from one of three independent experiments. (E) Fold increase in virions in culture supernatants at 4 dpi, as measured by RT activity in culture supernatants. Trans-complementation of the HIV-1_NL4-3_ IN_D116N_ virus with mutant IN molecules restored particle release to levels comparable to WT IN (red), partially restored particle release (gray) or could not restore particle release (blue). Columns show average of three independent experiments (open circles) and error bars represent standard deviation (*P < 0.05 and **P < 0.01, by paired t test between individual mutants and WT).

Three distinct phenotypes became apparent by assessing the amount of virion-associated IN and RT enzymes (Fig. 2A and Fig. S1A). First, virion-associated IN was at least 5-fold less than WT with several mutants, including H12N, N18I, K103E, W108R, F185K, L242A, and V260E (Fig. 2A and Table 1). Notably, these substitutions also reduced levels of Gag-Pol processing intermediates in producer cells (Fig. 2A) and RT in virions (Fig. 2A and Fig. S1A), suggesting that they likely destabilized the Gag-Pol precursor. Near complete lack of processing intermediates with the K14A and N18I substitutions, despite the presence of fully processed RT and IN in virions (detected using a separate polyclonal antibody), is likely due to inaccessibility of epitopes recognized by the monoclonal anti-IN antibody in the processing intermediates. Second, the R228A substitution abolished full-length IN in virions without impacting cell-or virion-associated Gag-Pol levels or processing intermediates; however, a faster migrating species due to altered charge in IN or that may represent product of aberrant IN processing and/or IN degradation was visible. A similar but more modest defect was observed for the K34A mutant, which was incorporated into virions at a modestly reduced level alongside a smaller protein species. Third, the remainder of the IN substitutions did not appear to affect IN or Gag-Pol levels in cells or virions.

**Table 1.** IN levels in virons.

| IN mutant | IN signal (%) | SD | IN mutant | IN signal (%) | SD |
| --- | --- | --- | --- | --- | --- |
| WT | 100 | N/A | K186E | 63.2 | 16.4 |
| H12N | ND | N/A | R187A | 34.4 | 24.5 |
| K14A | 40.5 | 10.3 | K188E | 45.8 | 26.3 |
| N18I | 16.7 | 3.1 | R199A | 41.1 | 19.2 |
| K34A | 32.6 | 6.9 | Q214L/Q216L | 31.7 | 6.3 |
| E87A | 31.4 | 16.1 | K215A/K219A | 47.7 | 20.4 |
| E96A | 38.4 | 31.2 | R228A | 50.2 | 26.4 |
| Y99A | 22.0 | 10.3 | K236E | 37.4 | 14.0 |
| K103E | 6.8 | 2.3 | L241A | 41.4 | 12.4 |
| W108R | 2.7 | 0.3 | L242A | 13.7 | 3.5 |
| V165A | 55.6 | 32.8 | K258A | 36.7 | 19.7 |
| F185K | 2.5 | 1.6 | V260E | 6.0 | 4.0 |
| K186A | 69.4 | 27.9 | R262A/R263A | 52.9 | 7.0 |

Quantitation of IN in virions as measured by immunoblotting. For each experiment IN signal was normalized to CA signal for each virus, and the resulting value compared to that of WT (set at 100%.) Reported values are the average value (as percent of WT) and standard deviation (SD) between 4 independent experiments. Mutants with less than 20% IN signal of WT are highlighted in gray.

With the exception of E96A, nearly all of the IN substitutions reduced virus titers at least 100-fold compared to the WT (Fig. 2B), which corresponded with reduced levels of reverse-transcription in infected cells (Fig. 2C). In line with previous reports [19, 20, 34], class II mutant IN molecules had variable levels of catalytic activity as assessed by the ability of Vpr-IN proteins to transcomplement a catalytically inactive IN (D116N, [11, 45]) in infected cells [68, 82]. All Vpr-IN fusion proteins, except for the H12N mutant which likely decreased the stability of the Vpr-IN fusion protein, were expressed at similar levels in cells (Fig. S1B). We found that K14A, E96A, Y99A, K103A, V165A, R187A, K188E R199A, K236E, and R269A/R273A IN mutants trans-complemented a catalytically inactive IN at levels similar to the WT, whereas W108R, R228A, and V260E mutants were unable to do so (Fig. 2D-E). The inability of W108R, R228A, and V260E mutants to transcomplement implies that they are impaired for integration, a result in line with previous observations [46, 81]. The remainder of the IN mutants restored integration, albeit at significantly lower than WT levels (Fig. 2D-E). These results suggest that the majority of the class II mutant INs retain structural integrity and at least partial catalytic activity in the presence of a complementing IN protein. Cumulatively, these data show that some class II substitutions in IN can affect the stability and/or processing of virion associated proteins, but they all universally lead to the formation of non-infectious virions that are blocked at reverse transcription in target cells, a hallmark of class II IN substitutions [10, 12].

### Class II IN mutants abolish IN binding to RNA

Using complementary in vitro and CLIP-based approaches, we have previously shown that IN interacts with the viral genome through multiple basic residues (i.e. K264, K266, R269, K273) in its CTD [28]. In addition, IN-RNA interactions could also depend on proper IN multimerization, as ALLINI-induced aberrant IN multimerization potently inhibited the ability of IN to bind RNA [28]. Based on this, in the next set of experiments, we aimed to determine whether class II IN mutants bind vRNA, and if not, whether improper IN multimerization may underlie this defect.

IN-vRNA complexes were immunoprecipitated from UV-crosslinked virions and the levels of coimmunoprecipitating vRNA was assessed. Note that substitutions that significantly reduced the amount of IN in virions (Fig. 2A, Table 1) were excluded from these experiments. All class II IN mutant viruses contained similar levels of vRNA, ruling out any inadvertent effects of the alterations on RNA packaging (Fig. 3A). While the catalytically inactive IN D116N bound vRNA at a level that was comparable to the WT, nearly all of the class II IN mutant proteins failed to bind vRNA (Fig. 3B). The E96A substitution, which had a fairly modest effect on virus titers as compared to other IN mutants (Fig. 2B), decreased but did not abolish the ability of IN to bind RNA (Fig. 3B). Thus, lack of RNA binding ability is a surprisingly common property of a disperse set of class II IN mutants, despite the fact that many of the altered amino acid residues are distally located from the CTD.

**Figure 3.**
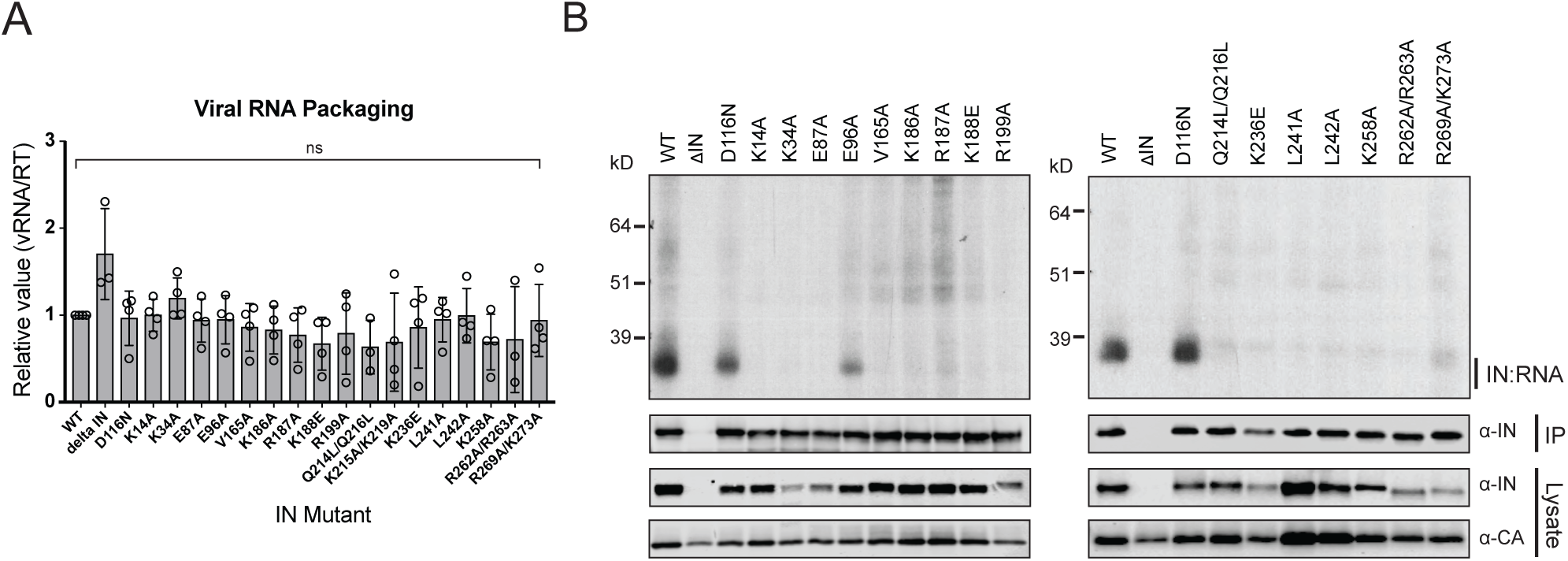
Class II IN substitutions prevent IN binding to the vRNA genome in virions. (A) Analysis of the levels of packaged viral genomic RNA in WT and IN mutant HIV-1_NL4-3_ virions. vRNA extracted from purified virions was measured by Q-PCR. Data was normalized to account for differences in particle yield using an RT activity assay. Normalized quantities of vRNA are expressed relative to WT (set to 1). Columns show the average of three-four independent experiments (open circles) and error bars represent standard deviation (ns, not significant, by one-way ANOVA). (B) Representative autoradiogram of IN-RNA adducts immunoprecipitated from WT or IN mutant HIV-1_NL4-3_ virions. The amount of immunoprecipitated material was normalized such that equivalent levels of WT and mutant IN proteins were loaded on the gel, as also evident in the immunoblots shown below. Levels of IN and CA in input virion lysates is shown in the lower immunoblots. Data is representative of three independent replicates.

### IN multimerization plays a key role in RNA binding

As it seemed unlikely that all of the class II IN substitutions directly inhibited IN binding to RNA, we reasoned that they might indirectly abolish binding by perturbing proper IN multimerization. To test whether class II IN substitutions altered IN multimerization in a relevant setting, purified HIV-1_NL4-3_ virions were treated with ethylene glycol bis (succinimidyl succinate) (EGS) to covalently crosslink IN in situ and virus lysates were analyzed by immunoblotting. IN species that migrated at molecular weights consistent with those of monomers, dimers, trimers and tetramers were readily distinguished in WT virions (Fig. 4A). In the majority of the class II mutant particles, IN appeared to be predominantly monomeric, with little dimers and no readily detectable tetramers (Fig. 4A). In contrast to this general pattern, K34A, E96A, R262A/R263A and R269A/K273A IN mutants formed dimers and tetramers at similar levels to the WT (Fig. 4A). An undefined smear was present at higher molecular weights for all virions, possibly as a result of the formation of large IN aggregates upon cross-linking (Fig. 4A).

**Figure 4.**
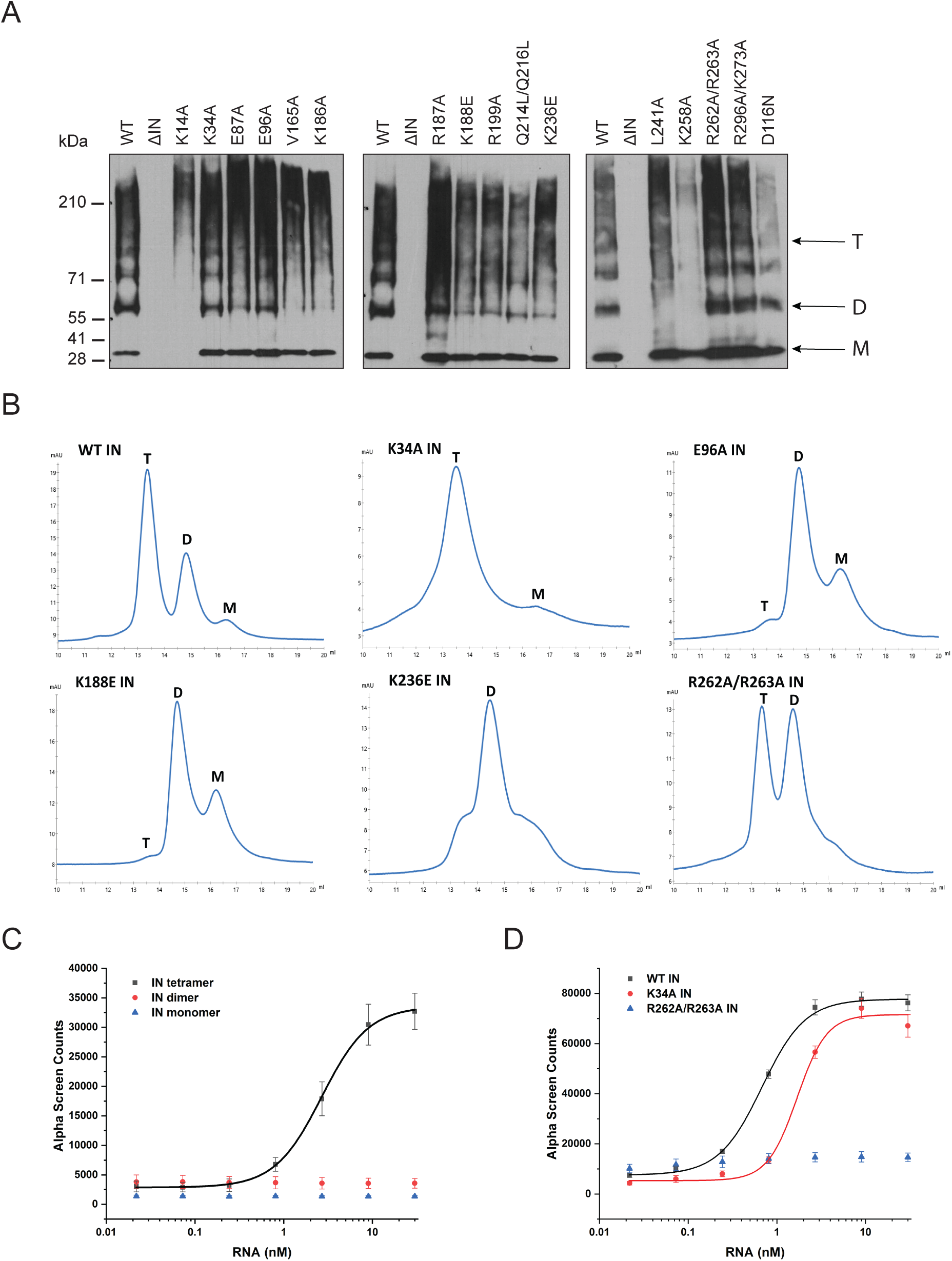
Multimerization properties of class II IN mutants. (A) Immunoblot analyses of IN multimers in virions. Purified WT or IN mutant HIV-1_NL4-3_ virions were treated with 1 mM EGS, and virus lysates analyzed by immunoblotting using antibodies against IN following separation on 6% Tris-acetate gels. Position of monomers (M), dimers (D), and tetramers (T) are indicated by arrows. Representative image of one of three independent experiments is shown. (B) SEC profiles of 10 µM INs by Superdex 200 10/300 GL column. X-axis indicates elution volume (mL) and Y-axis indicate the intensity of absorbance (mAU). Tetramers (T), Dimers (D) and Monomers (M) are indicated. Representative chromatograms from two independent analyses are shown. (C) Analysis by Alpha screen assay of 100 nM WT IN monomers, dimers, and tetramers binding to biotinylated TAR RNA after separation by SEC. Graphed data is the average of three independent experiments and error bars indicate standard deviation. (D) Analysis of 100 nM WT or mutant INs binding to biotinylated TAR RNA by Alpha screen assay. Graphed data is the average of three independent experiments and error bars indicate standard deviation.

To corroborate these findings, we analyzed the multimerization properties of recombinant WT, K34A, E96A, K188E, K236A and R262A/R263A IN proteins by SEC (Fig. 4B). In line with the crosslinking studies in virions, WT, K34A and R262A/R263A IN molecules all formed tetramers, while the levels of dimers varied between the mutants. For example, while IN R262A/R263A presented similar levels of tetramers and dimers, IN K34A was primarily tetrameric with a minor dimeric species, as evident by the broad right shoulder of the tetrameric SEC peak (Fig. 4B). In contrast, E96A and K188E IN molecules almost exclusively formed dimers and monomers with little evidence for tetramer formation (Fig. 4B). While K236E IN was predominantly dimeric, the broad base of its chromatogram revealed some evidence for tetramers and monomers as well (Fig. 4B).

We next determined the ability of IN monomers, dimers and tetramers to bind RNA to test whether there is a causal link between the multimerization defects of class II IN substitutions and RNA binding. Following size-exclusion chromatography (SEC)-based separation of monomeric, dimeric and tetrameric IN, their affinity for TAR RNA, which constitutes a high affinity binding site for IN [28], was assessed by an Alpha-screen assay. Remarkably, while WT IN tetramers bound to TAR RNA at high affinity (2.68 ± 0.16 nM), neither IN dimers nor monomers showed evidence of binding (Fig. 4C). Although IN K34A and IN R262A/R263A could both form tetramers, IN K34A showed a reduced affinity for RNA while IN R262A/R263A could not bind RNA at all (Fig. 4D).

Collectively, these results pointed to a key role of IN tetramerization in RNA binding and suggest that a defect in proper multimerization underlies the inability of the majority of class II IN mutants to bind vRNA. As the K34A and R262A/R263A substitutions did not affect IN tetramerization, our findings suggest that these residues may be directly involved in IN binding to RNA.

### Class II IN substitutions generate virions with eccentric morphology

We next sought to determine how preclusion or inhibition of IN-vRNA interactions correlated with particle morphology. Virion morphology of a subset of the IN mutants that inhibited vRNA interactions by three different mechanisms; i.e. those that decreased IN levels in virions (N18I and W108R), those that may have directly inhibited IN binding to RNA (K34A, R262A/R263A), and those that primarily altered IN multimerization (E87A, E96A, F185K, R187A, L241A, L242A), was assessed by transmission electron microscopy (TEM). As expected, the majority of WT particles contained an electron dense condensate representing vRNPs inside the CA lattice, whereas an ΔRT-IN deletion mutant virus produced similar levels of immature particles and eccentric particles (Fig. 5A-B). Remarkably, irrespective of how IN-RNA interactions were inhibited, 70-80% of nearly all class II IN mutant particles exhibited an eccentric morphology (Fig. 5A-B). Of note, the E96A mutant tended to produce less eccentric and more mature particles than the other IN mutants. Because IN E96A retained partial binding to vRNA in virions (Fig. 3B) and partial infectivity (Fig. 2B), we conclude that this infection-deferred mutant harbors a partial class II phenotype.

**Figure 5.**
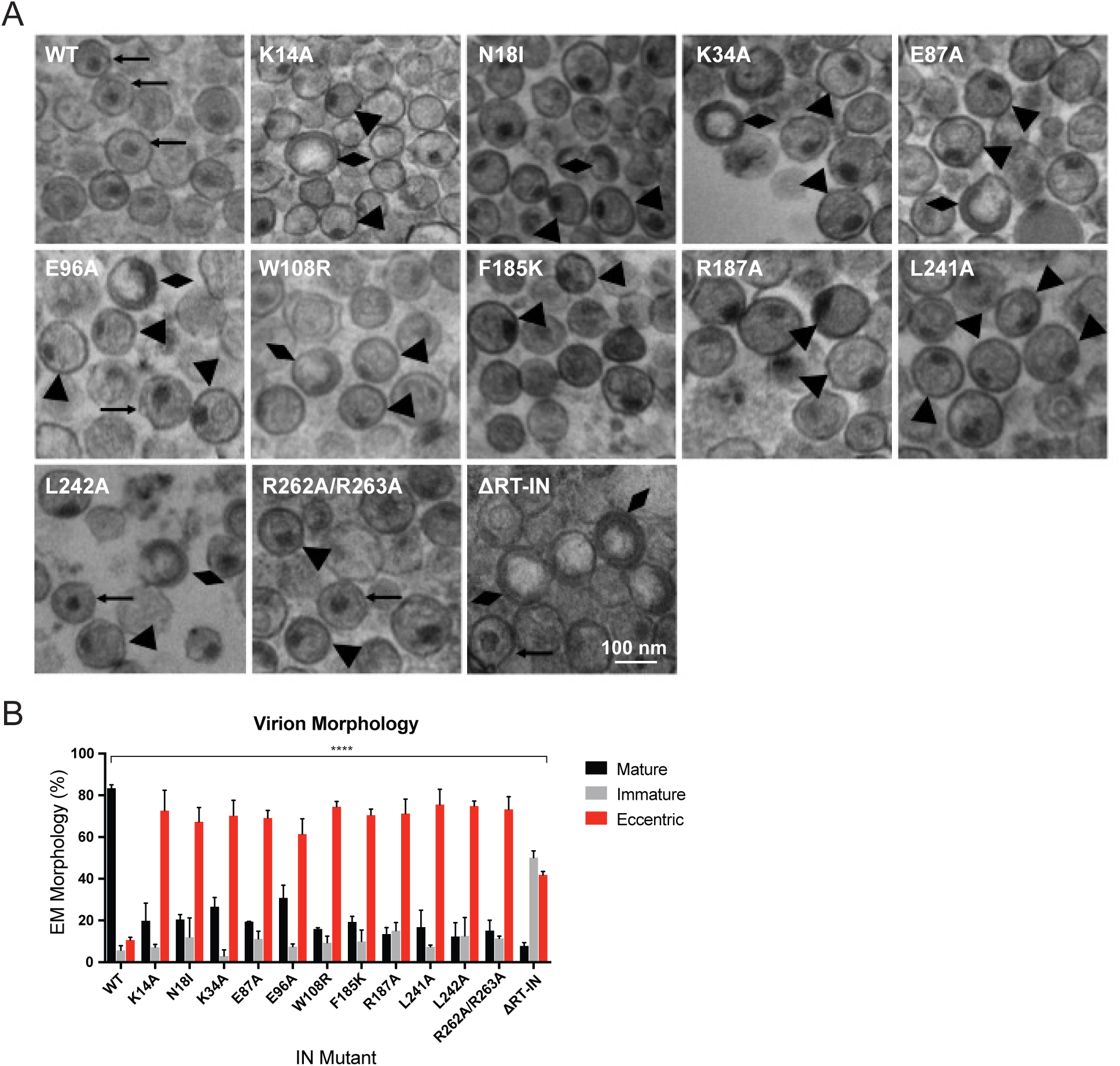
Analysis of class II IN mutant virion morphologies viruses by TEM. (A) Representative TEM images of WT, K14A, N18I, K34A, E87A, E96A, W108R, F185K, R187A, L241A, L242A, R262A/R263A, and ΔRT-IN HIV-1_NL4-3_ virions. Magnification is 30,000x (scale bar, 100 nm). Black arrows indicate mature particles containing conical or round cores with associated electron density; triangles indicate eccentric particles with electron dense material situated between translucent cores and the viral membrane; diamonds indicate immature particles. (B) Quantification of virion morphologies. Columns show the average of two independent experiments (more than 100 particles counted per experiment) and error bars represent standard deviation.

Next, we tested whether inhibition of IN-RNA interactions through class II substitutions changes the localization of IN in virions. The premise for this is based on our previous finding that disruption of IN binding to vRNA through the IN R269A/K273A substitution leads to separation of a fraction of IN from dense vRNPs and CA containing complexes [64]. Thus, we predicted that inhibition of IN-RNA interactions through the above class II substitutions could lead to a similar outcome. To this end, WT or class II IN mutant virions stripped of the viral lipid envelope by brief detergent treatment were separated on sucrose gradients, and resulting fractions were analyzed for CA, IN, and matrix (MA) content by immunoblotting [64, 83]. As before [64], WT IN migrated primarily in dense fractions, whereas the R269A/K273A mutant migrated bimodally (Fig. 6A, B). In contrast to our hypothesis, the majority of IN mutants sedimented similarly to WT IN and settled in the denser gradient fractions (Fig. 6A, B). Exceptions were the K34A and R262A/R263A IN mutants, a fraction of which migrated in soluble fractions similar to the R269A/K273A mutant, suggesting their localization outside of the capsid lattice. None of the IN substitutions affected the migration pattern of CA (Fig. 6C), which distributed bimodally between the soluble and dense fractions, nor the distribution of MA (data not shown), which was found in mainly the soluble fractions. These results suggested that, with the exception of the K34A, R262A/R263A, and R269A/K273A, IN mutant proteins may remain associated with the CA lattice despite inhibition of IN-vRNA interactions.

**Figure 6.**
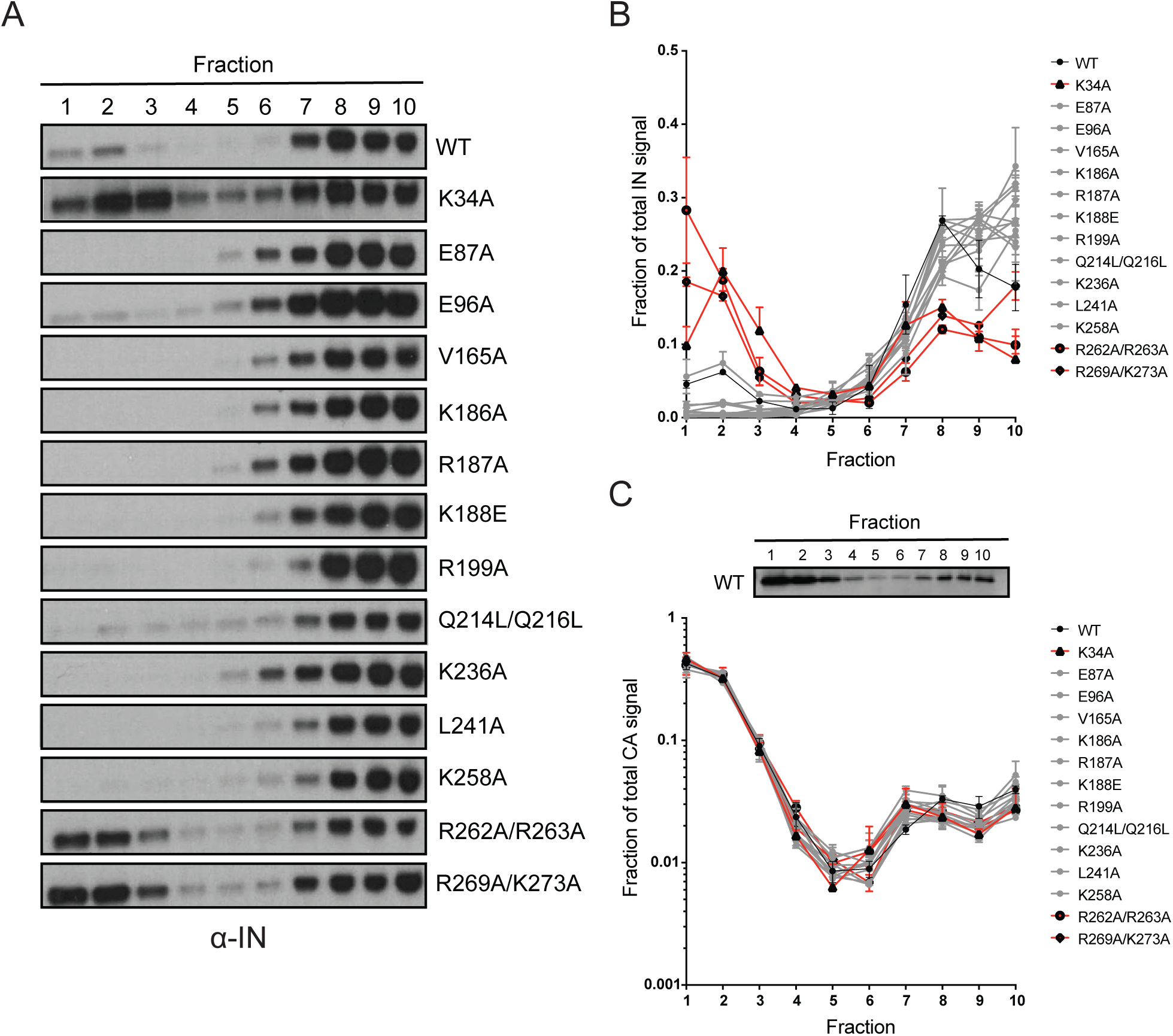
Biochemical analysis of class II IN mutant virus particles. (A) Immunoblot analysis of sedimentation profiles of IN in WT or IN mutant virions. Purified HIV-1_NLGP_ virions were analyzed by equilibrium density centrifugation as detailed in Materials and Methods. Ten fractions collected from the top of the gradients were analyzed by immunoblotting using antibodies against IN. Representative images from one of four independent experiments are shown. (B) Quantitation of IN signal intensity in immunoblots as in (A) are shown. Profile of WT virions is denoted in black, IN mutants that led to bimodal IN distribution are shown in red and others are shown in grey. Graphed data is the average of two independent experiments and error bars indicate the range. (C) Representative immunoblot analysis of sedimentation profile of CA in WT virions and quantitation of CA signal intensity in immunoblots are shown. Profile of WT virions is denoted in black, IN mutants that led to bimodal IN distribution are shown in red and others are shown in grey. Graphed data is the average of two independent experiments and error bars indicate the range.

### Premature loss of vRNA and IN from class II IN mutant viruses upon infection of target cells

We have previously shown that vRNA and IN are prematurely lost from cells infected with the R269A/K273A class II IN mutant [64]. Given that eccentric vRNP localization is a common feature of class II IN mutant viruses (Fig. 5), we next asked whether loss of vRNA in target cells is a common outcome for other class II IN mutant viruses. As the majority of mutant IN molecules appeared to remain associated with higher-order CA in virions (Fig. 6), we also wanted to test whether they would be protected from premature degradation in infected cells.

The fates of viral core components in target cells were tracked using a previously described biochemical assay [74]. For these experiments we utilized pgsA-745 cells (pgsA), which lack surface glycosaminoglycans, and likely as a result can be very efficiently infected by VSV-G-pseudotyped viruses in a synchronized fashion. PgsA cells were infected with WT or IN mutant viruses bearing substitutions that inhibited IN-vRNA interactions directly and led to mislocalization of IN in virions (i.e. K34A, R262A/R263A, R269A/K273A) or indirectly through aberrant IN multimerization and did not appear to grossly affect IN localization in virions (i.e. E87A, V165A) (Fig 7A). Following infection, post-nuclear lysates were separated on linear sucrose gradients, and fractions collected from gradients were analyzed for viral proteins (CA, IN, RT) and vRNA by immunoblotting and Q-PCR-based assays, respectively.

**Figure 7.**
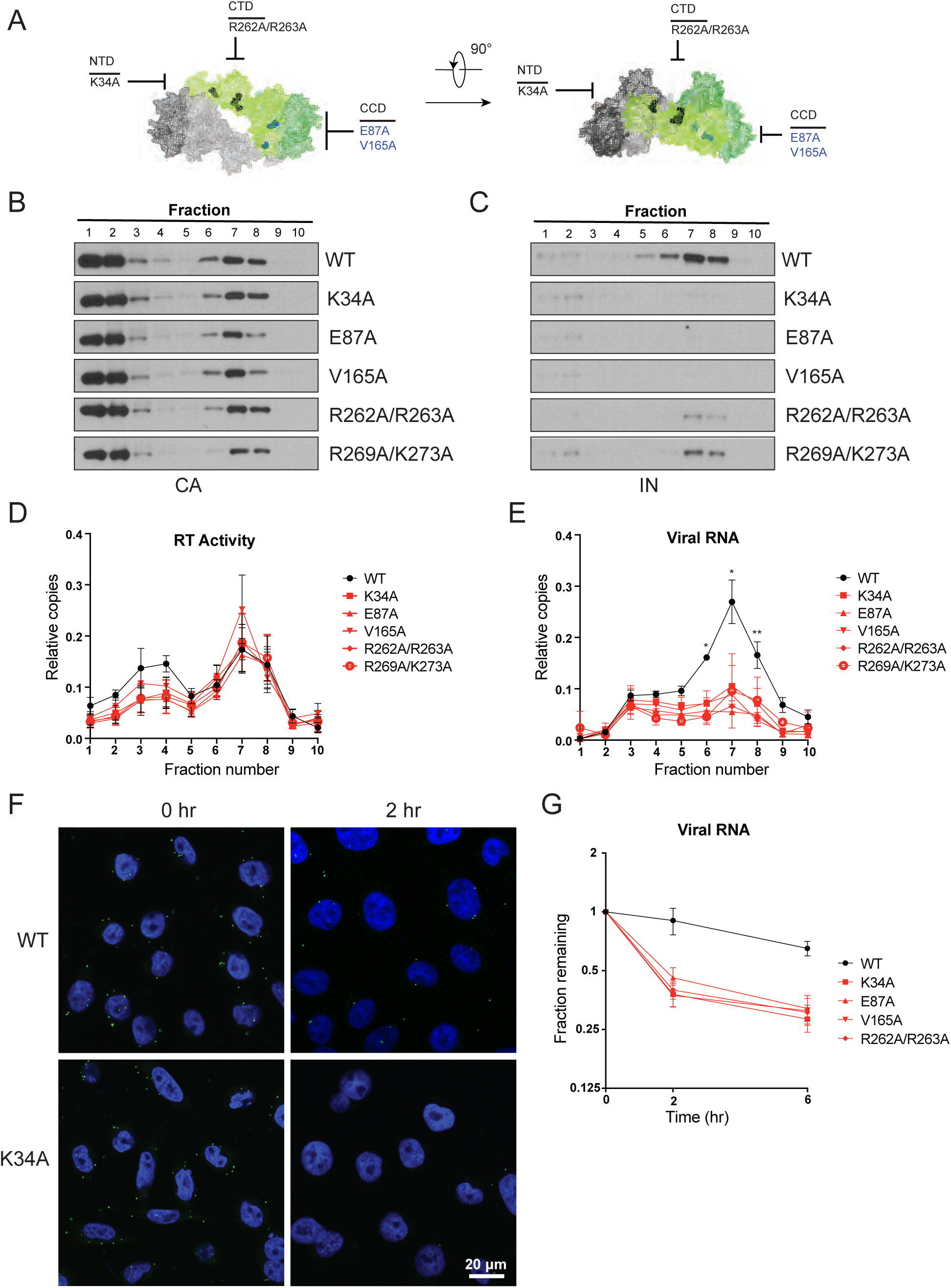
Premature loss of vRNA and IN from class II IN mutant viruses upon infection of target cells. (A) Locations of the class II IN substitutions K34A, E87A, V165A and R262A/R263A displayed on a single IN monomer within the context of the HIV-1 IN tetramer intasome structure (PDB 5U1C.) Substitutions are color coded based on whether they putatively caused mislocalization of IN in virions (black) or not (blue.) (B-E) PgsA-745 cells were infected with WT or IN mutant HIV-1 virions and fates of viral core components were analyzed 2 hpi. Fractions were analyzed for the presence of CA (B) and IN (C) by immunoblotting and for RT activity (D) and vRNA (E) by Q-PCR. Immunoblots are representative of three independent experiments. Graphed data in (D) and (E) is the average of three independent experiments with error bars indicating standard deviation (*P < 0.05 and **P < 0.01, by repeated measures one-way ANOVA.) (F) Representative images of pgsA-745 cells infected with WT or IN mutant HIV-1_NL4-3_ viruses 0 and 2 hpi. Cells were stained for vRNA (green) and nuclei (blue) as detailed in Materials and Methods. (G) Fraction of viral RNA remaining after 2 and 6 hpi compared to the quantity measured at 0 hpi. MT-4 cells were synchronously infected with VSV-G pseudotyped HIV-1_NL4-3_ viruses and at each timepoint samples of infected cultures were taken for analysis. Viral RNA levels in samples were measured by Q-PCR and normalized to the levels of GAPDH mRNA. Data points are the average of five independent experiments with error bars indicating standard error of the mean.

As previously reported [64, 74], in cells infected with WT viruses, IN, RT, vRNA and a fraction of CA comigrated in sucrose fractions 6-8, representing active RTCs (Fig. 7B-E). Note that a large fraction of CA migrated in the top two soluble sucrose fractions representing CA that had dissociated from the core as a result of uncoating or CA that was packaged into virions but not incorporated into the capsid lattice [84, 85]. Notably, in cells infected with class II IN mutant viruses, equivalent levels of CA (Fig. 7B) and RT (Fig. 7D) remained in the denser fractions, whereas IN (Fig. 7C) and vRNA (Fig. 7E) were substantially reduced. Loss of vRNA and IN from dense fractions, without any corresponding increase in the top fractions containing soluble proteins and RNA, suggest their premature degradation and/or mislocalization in infected cells.

We next employed a complementary microcopy-based assay [75] in the context of full-length viruses to corroborate these findings. Advantages of this approach over biochemical fractionation experiments include the ability to track HIV-1 vRNA at the single-cell level with a high degree of specificity (Fig. S2A), determine its subcellular localization, and to side-step possible post-processing artifacts associated with biochemical fractionation. Cells were synchronously infected with VSV-G pseudotyped HIV-1_NL4-3_ in the presence of nevirapine to prevent vRNA loss due to reverse transcription, and vRNA levels associated with cells immediately following synchronization (0 h) and 2 h post-infection were evaluated [75]. In WT-infected cells, vRNA was clearly visible immediately after infection (Fig. 7F). Two h post infection, cell associated vRNA had fallen to 60-80% of starting levels (Fig. 7F, S2C), likely as the result of some viruses failing to enter or perhaps being degraded after entry. However, a significant proportion of vRNA was still readily detectable. In contrast, in cells infected with the IN mutant viruses the reduction in vRNA was greater, and by 2 h post-infection only 30-40% remained (Fig. 7F and Fig. S2B-C). These results support the conclusion from the biochemical fractionation experiments that vRNA is prematurely lost from cells infected with class II IN mutant viruses.

Finally, we tested whether our findings held true in physiologically relevant human cells. MT-4 T cells were synchronously infected with WT or class II IN mutant VSV-G pseudotyped HIV-1_NL4-3_ in the presence of nevirapine. Cells were collected immediately after synchronization (0 h), 2 and 6 h post-infection, and the quantity of vRNA measured by Q-PCR. In line with the above findings, vRNA levels decreased at a faster rate with the class II IN mutants as compared to WT viruses, with half as much cell-associated vRNA remaining at 2 and 6 h post-infection for the class II IN mutants (Fig. 7G). Treating cells with ammonium chloride to prevent fusion of the VSV-G pseudotyped viruses rescued vRNA loss, and vRNA from WT and mutant viruses were retained at equal levels, indicating that the loss of vRNA is dependent on entry into the target cell (Fig. S2D). These findings agree with the previous experiments and demonstrate that class II IN substitutions lead to the premature loss of vRNA genome also in human T cells.

## DISCUSSION

Our findings highlight the critical role of IN-vRNA interactions in virion morphogenesis and provide the mechanistic basis for how diverse class II IN substitutions lead to similar morphological and reverse transcription defects. We propose that class II IN substitutions lead to the formation of eccentric particles through three distinct mechanisms (Fig. 8): (i) depletion of IN from virions thus precluding the formation of IN-vRNA complexes; ii) impairment of functional IN multimerization and as a result, indirect disruption of IN-vRNA binding; iii) direct disruption of IN-vRNA binding without substantially affecting IN levels or its inherent multimerization properties. Irrespective of how IN binding to vRNA is inhibited, all substitutions led to the formation of eccentric viruses that were subsequently blocked at reverse transcription in target cells. We provide evidence that premature degradation of the exposed vRNPs and separation of RT from the vRNPs underlies the reverse transcription defect of class II IN mutants (Fig. 7). Taken together, our findings cement the view that IN binding to RNA accounts for the role of IN in accurate particle maturation and provide the mechanistic basis of why these viruses are blocked at reverse transcription in target cells.

**Figure 8.**
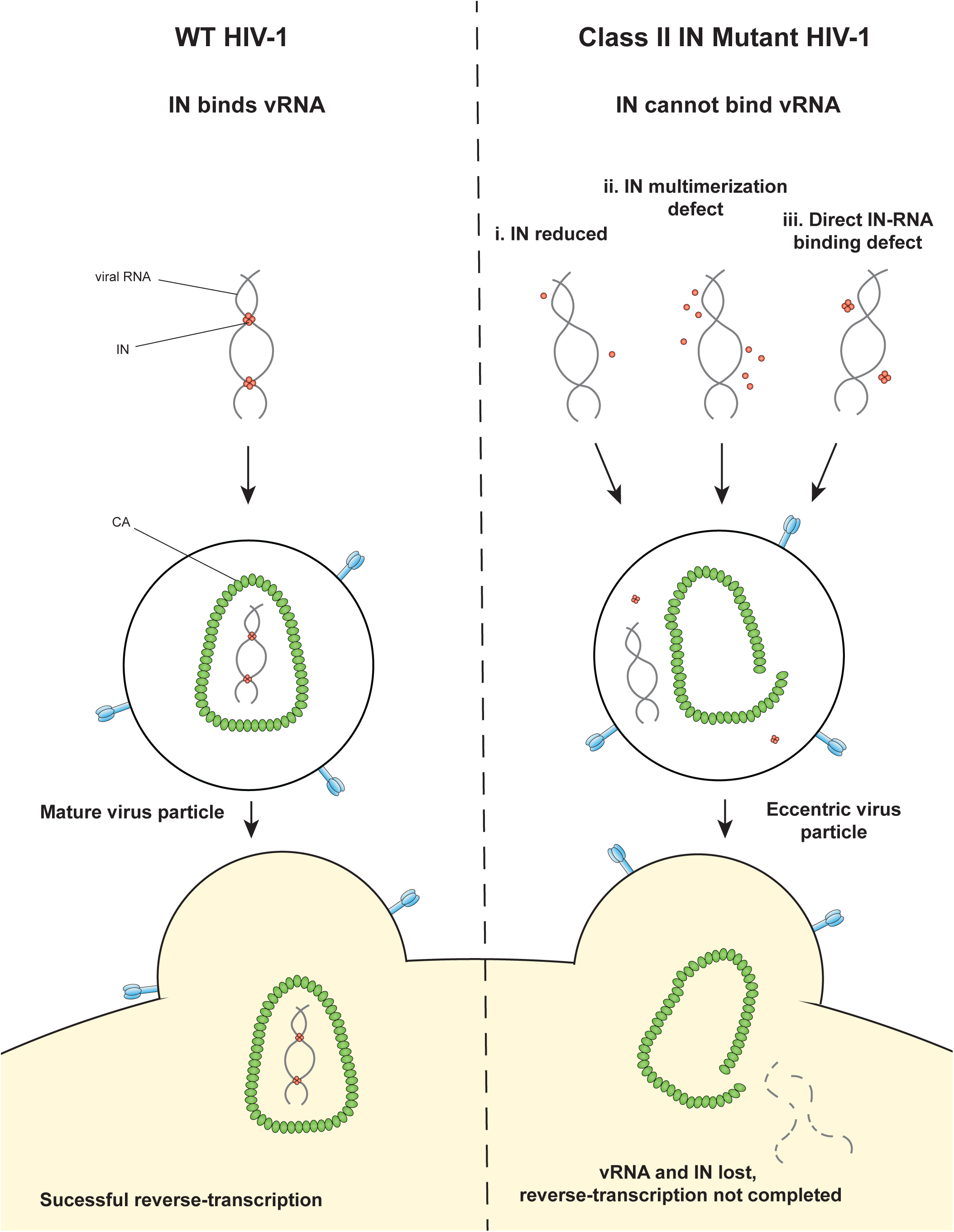
Model depicting how class II IN mutants exert their effects on HIV-1 replication.

In regard to the first case (i) above, it was previously shown that IN deletion leads to the formation of eccentric particles [11, 27]. Thus, it is reasonable to assume that missense mutations that decreased IN levels in virions phenocopy IN deletion viruses. While it is also possible that these substitutions additionally affected IN binding to vRNA or multimerization, we could not reliably address these possibilities due to the extremely low levels of these proteins in virions.

Our results show the striking affinity of IN tetramers to bind RNA compared with IN monomers and dimers (Fig. 4C). In support of tetramerization being a prerequisite for RNA-binding, the inability of a number of class II IN mutant proteins to bind RNA was accompanied by a clear multimerization defect both in virions (Fig. 4A) and in vitro (Fig. 4B). The structural basis for IN binding to RNA is not yet known; however, these findings are in line with the previous in vitro evidence that hinted a link between IN multimerization and RNA-binding. For example, IN binds RNA as lower-order multimers, and conversely RNA binding may inhibit the formation of higher order IN multimers in vitro [28]. Notably, formation of open IN polymers that occlude the IN CTD from RNA binding may underlie the inhibition of IN-RNA interactions by ALLINIs [28, 58].

Based on MS-based footprinting experiments in vitro, we previously found that positively charged residues within the CTD of IN (i.e. K264, K266, K273) directly contact RNA, as was also validated by CLIP experiments [28]. Our findings here suggest that IN-vRNA contacts may extend to nearby basic residues within the CTD, such as R262 and R263, and perhaps more surprisingly, K34 within the IN NTD, as alterations of these residues did not prevent IN tetramerization (Fig. 4A-B) but completely abolished IN-vRNA binding in virions (Fig. 3B) and reduced RNA-binding in vitro (Fig 4D). This raises the possibility of a second RNA-binding site in the IN NTD. Structural analysis of IN in complex with RNA will be essential to definitively determine how IN binds RNA as well as the precise multimeric species required for binding.

The mechanism by which IN-vRNA interactions mediate the encapsidation of vRNPs inside the CA lattice remains unknown. One possibility is that the temporal coordination of proteolytic cleavage events during maturation is influenced by IN-vRNA interactions [86, 87]. In this scenario, the assembly of the CA lattice may become out of sync with the compaction of vRNA by NC. Another possibility is that IN-vRNA complexes nucleate the assembly of the CA lattice, perhaps by directly binding to CA. Notably, the biochemical assays performed herein show that class II IN substitutions do not appear to affect the assembly and stability of the CA lattice in vitro and in target cells. Although this finding is in disagreement with the previously observed morphological aberrations of the CA lattice present in eccentric particles [26], it is possible that the biochemical experiments used herein lack the level of sensitivity required to quantitatively assess these aberrations. Further studies deciphering the crosstalk between IN-RNA interactions and CA assembly will be critical to our understanding of the role of IN in accurate virion maturation.

While the mislocalization of the vRNA genome in eccentric particles can be accurately assessed by TEM analysis, precisely where IN is located in eccentric particles remains an open question. Earlier studies based on biochemical separation of core components from detergent-treated IN R269A/K273A virions indicated that IN may also mislocalize outside the CA lattice [64]. In this study, only two class II IN mutants (K34A and R262A/R263A) revealed this phenotype (Fig. 6A, B). It is intriguing that the bimodal distribution of IN in this experimental setting was only seen with IN mutants that directly inhibited IN binding to vRNA. A possible explanation for these observations is that improperly multimerized IN is retained within the CA lattice or in association with it. Despite this co-migration pattern with CA, we found that both E87A and V165A mutant INs were rapidly lost in infected cells, suggesting that they are not fully protected by CA upon cellular entry (Fig. 7C).

Why is the unprotected vRNA and IN prematurely lost in target cells? It seems evident that the protection afforded by the CA lattice matters the most for vRNP stability, though we cannot rule out that IN binding to vRNA may in and of itself stabilize both the genome and IN. Alternatively, the AU-rich nucleotide content of HIV-1 may destabilize its RNA [88–90], similar to several cellular mRNAs that encode for cytokines and growth factors [91]. Finally, RNA nicking and deadenylation in virions by virion associated enzymes [92–94] may predispose retroviral genomes to degradation when they are prematurely exposed to the cytosolic milieu. While cytosolic IN undergoes proteasomal degradation when expressed alone in cells [95–99], we have found previously that proteasome inhibition does not rescue vRNA or IN in target cells infected with a class II IN mutant [64]. Further studies are needed to determine whether a specific cellular mechanism or an inherent instability of vRNPs is responsible from the loss of vRNA in infected cells.

In conclusion, we have identified IN-vRNA binding as the underlying factor for the role of IN in virion morphogenesis and show that virion morphogenesis is necessary to prevent the premature loss of vRNA and IN early in the HIV-1 lifecycle. Despite relatively high barriers, drugs that inhibit the catalytic activity of IN do select for resistance, and additional drug classes that inhibit IN activity through novel mechanisms of action would be a valuable addition to currently available treatments. The finding that IN-vRNA interaction can be inhibited in multiple ways-by directly altering residues in the IN CTD or by altering IN multimerization in virions-can help guide the design of future anti-retroviral compounds.

---

**Figure S1.**
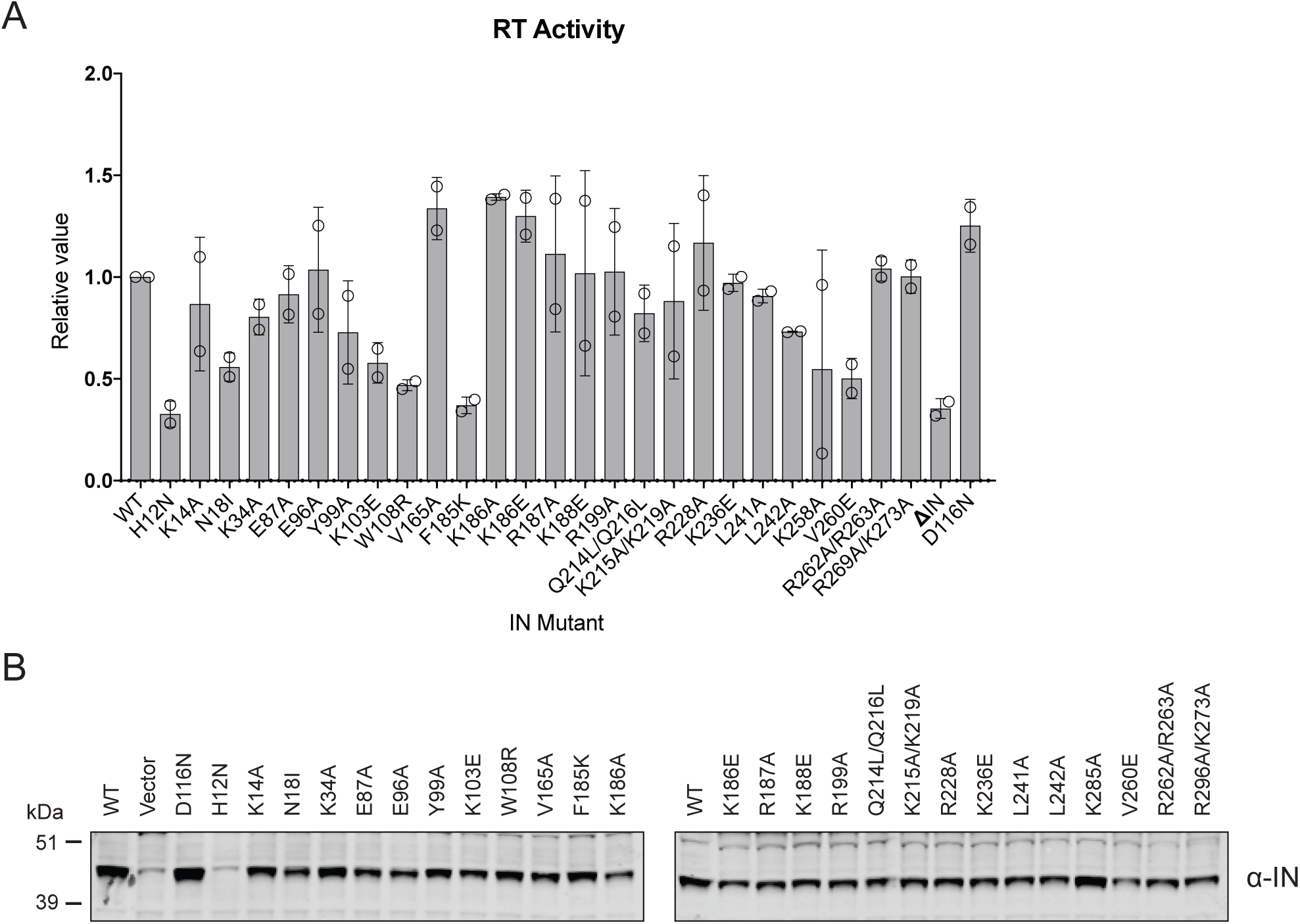
Characterization of the replication defects of class II IN mutant viruses. (A) Reverse-transcriptase activity measured in HIV-1_NL4-3_ virion lysates. For each repetition RT activities for the IN mutants are expressed relative to the WT (set to 1.) Columns show average of two independent experiments (open circles) and error bars represent standard deviation (****P < 0.0001, ***P < 0.001, **P < 0.01, and *P < 0.05, by unpaired t test between individual mutants and WT.) (B) Representative immunoblot analysis of Vpr-IN fusion constructs in cell lysates. HEK293T cells were co-transfected with the HIV-1_NL4-3_ IN_D116N_ proviral plasmid along with Vpr-IN expression plasmids encoding for the indicated IN substitutions or an empty vector control. Expression of Vpr-IN constructs in cell lysates was detected using an anti-IN antibody.

**Figure S2.**
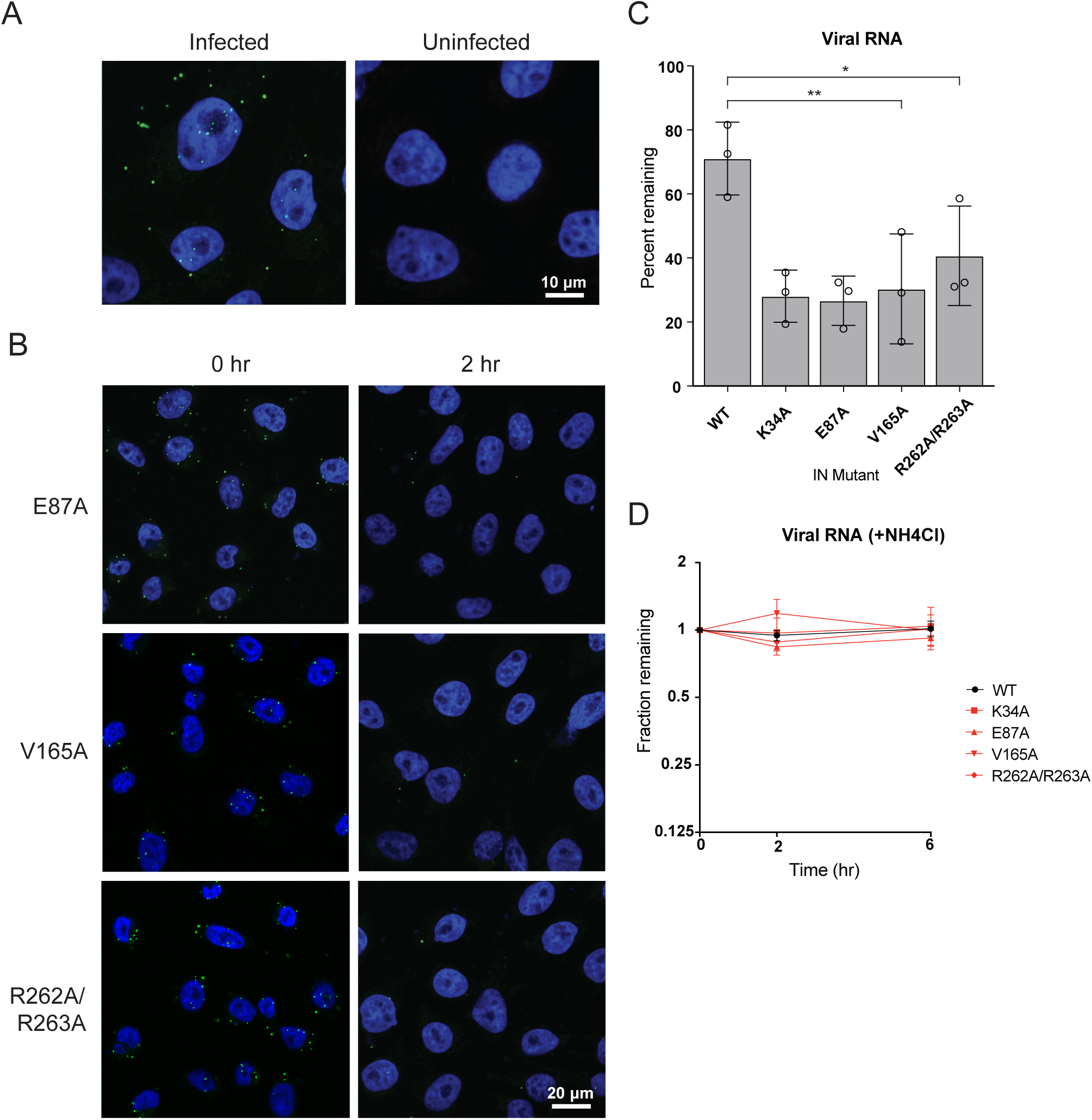
Premature loss of vRNA and IN from class II IN mutant viruses upon infection of target cells. (A) Representative images of uninfected pgsA-745 cells and cells infected with WT HIV-1_NL4-3_ viruses at 0 hpi. Cells were fixed and stained for vRNA (green) and nuclei (blue). (B) Representative images of pgsA745 cells infected with IN mutant HIV-1_NL4-3_ viruses 0 and 2 hpi. Cells were fixed and stained for vRNA (green) and nuclei (blue). (C) Quantification of vRNA remaining in cells infected with WT or IN mutant HIV-1_NL4-3_ viruses at 2 hpi. Values are the percent of vRNA remaining at 2 hpi compared to at 0 hpi. Columns show average of three independent experiments (open circles) and error bars represent standard deviation (*P < 0.05 and **P < 0.01, by one-way ANOVA with Dunnett’s multiple comparison test.) (D) Fraction of viral RNA remaining after 2 and 6 hpi compared to the quantity measured at 0 hpi. MT-4 cells were synchronously infected with VSV-G pseudotyped HIV-1_NL4-3_ viruses and incubated in the presence of 50 mM ammonium chloride for 6 hrs. At each timepoint samples of infected cultures were taken for analysis and levels of viral RNA in samples were measured by Q-PCR and normalized to the levels of GAPDH mRNA. Data points are the average of three independent experiments with error bars indicating standard error of the mean.

## ACKNOWLEDGEMENTS

We thank Dr. Michael Malim for providing the anti-IN monoclonal antibody and members of the Tolia lab for assisting in PyMol analysis. This study was supported by grants NIH grants P50 GM103297 (the Center for HIV RNA Studies) and GM122458 to SBK, AI143389-F31 fellowship to JE, R01 AI062520 to MK and SBK, U54 AI150472 to MK and AE, AI070042 to AE.

